# Discovery and Development of Human SARS-CoV-2 Neutralizing Antibodies using an Unbiased Phage Display Library Approach

**DOI:** 10.1101/2020.09.27.316174

**Authors:** Xia Cao, Junki Maruyama, Heyue Zhou, Lisa Kerwin, Rachel Sattler, John T. Manning, Sachi Johnson, Susan Richards, Yan Li, Weiqun Shen, Benjamin Blair, Na Du, Kyndal Morais, Kate Lawrence, Lucy Lu, Chin-I Pai, Donghui Li, Mark Brunswick, Yanliang Zhang, Henry Ji, Slobodan Paessler, Robert D. Allen

## Abstract

SARS-CoV-2 neutralizing antibodies represent an important component of the ongoing search for effective treatment of and protection against COVID-19. We report here on the use of a naïve phage display antibody library to identify a panel of fully human SARS-CoV-2 neutralizing antibodies. Following functional profiling *in vitro* against an early pandemic isolate as well as a recently emerged isolate bearing the D614G Spike mutation, the clinical candidate antibody, STI-1499, and the affinity-engineered variant, STI-2020, were evaluated for *in vivo* efficacy in the Syrian golden hamster model of COVID-19. Both antibodies demonstrated potent protection against the pathogenic effects of the disease and a dose-dependent reduction of virus load in the lungs, reaching undetectable levels following a single dose of 500 micrograms of STI-2020. These data support continued development of these antibodies as therapeutics against COVID-19 and future use of this approach to address novel emerging pandemic disease threats.

## INTRODUCTION

Global incidence of severe acute respiratory disease syndrome coronavirus 2 (SARS-CoV-2) infection has continued to rapidly increase since the virus was first detected in December 2019 ^1^. To date, public health agency efforts to combat the pandemic level of infection and resultant Coronavirus disease 2019 (COVID-19) have relied mostly upon effective quarantine measures ^2, 3^. To further protect at-risk populations, a worldwide effort has been undertaken to develop additional countermeasures, such as antiviral compounds, immunomodulatory agents, vaccines, and neutralizing antibodies (nAbs) ^4, 5, 6^.

While vaccines have been among the most effective means in preventing and containing widespread infections caused by many pathogens, their development is often faced with drawbacks, including long development timelines, large clinical trial sizes, and uncertain efficacy due to the reliance on patient-generated immune responses to the antigens^7, 8, 9^. Of note, it has been shown in the context of multiple virus infections that virus-specific antibodies can lead to exacerbation of disease symptoms through a process termed antibody dependent enhancement (ADE) ^10, 11^. This ADE of virus infection is a phenomenon in which certain virus-specific antibodies enhance the entry of virus due to engagement between IgG Fc regions and Fc-receptors on the surface of monocytes leading to productive infection of these cells rather than virus clearance ^12, 13^. In addition, the presence of immunodominant epitopes, i.e. highly immunogenic sequences, in viral antigens can skew this immune response, resulting in production of antibodies that are neither neutralizing nor protective, as seen in the vaccine development against human immunodeficiency virus (HIV)^14^. In the case of SARS-CoV-2, nAbs targeting the Spike protein have been isolated from the sera of convalescent COVID-19 patients as well as from sera donated by healthy normal patients prior to December 2019 ^15, 16, 17, 18, 19, 20^. While the patients’ immune systems generated these nAbs, they often recognize the same region of the Spike protein or even a similar epitope due to immunodominant regions of the Spike protein ^21, 22^. One way to overcome this inherent immune bias is the use of an artificial immune system in a test tube, namely phage display human antibody libraries, which allows for functional unbiased selection of Spike-binding antibodies irrespective of the underlying natural immunogenicity ^23^. Most antibody libraries are constructed in a way that allows for chain shuffling leading to random associations of heavy and light chain encoding genes and resulting in a vast repertoire of non-naturally occurring antigen specificities. In fact, few candidate anti-SARS-CoV-2 nAbs have been isolated from phage display libraries ^24, 25^. Using Sorrento’s G-MAB™ library ^26^, a single chain variable fragment (scFv) antibody phage display library constructed from the antibody repertoire of over 600 healthy individuals, a panel of antibodies that bind the SARS-CoV-2 Spike S1 subunit was identified and characterized.

Herein, we report on the identification, characterization, and subsequent optimization of STI-1499, a SARS-CoV-2 neutralizing antibody isolated from the G-MAB library. STI-1499 neutralized both the WA-1/2020 isolate of SARS-CoV-2 as well as isolate 2020001, a clinical isolate of SARS-CoV-2 that encodes the G614 variant of the Spike protein. Isolates harboring the G614 Spike variant have been reported to be more infectious and occur in greater frequency than viruses such as the WA-1/2020 isolate that encode the D614 Spike variant^27^. Affinity maturation of STI-1499 resulted in identification of STI-2020, an antibody with a 35-fold increased affinity for the SARS-CoV-2 Spike receptor-binding domain (RBD) leading to a greater than 50-fold increase in virus neutralization potency against live WA-1/2020 and 2020001 viruses *in vitro*. Importantly, both nAbs, STI-1499 and STI-2020, provided protection against the pathogenic effects and replication of SARS-CoV-2 in the Syrian golden hamster virus challenge model ^28, 29^.

## MATERIALS AND METHODS

### Antibody binding ELISA

The S1 subunit of the SARS-CoV-2 spike protein (amino acids 16-685) bearing a C-terminal histidine tag (ACRO Biosystems, Newark, NJ) was coated at 2 μg/ml on a Ni-NTA plate (Qiagen, Valencia, CA). After washing and blocking, 10 μg/ml antibody was added to the corresponding wells for 1-hour incubation. The plate was washed three times and incubated with 1:5000 dilution of HRP-conjugated mouse anti-human IgG (Fc). Color development was performed with 3,3′,5,5′-Tetramethylbenzidine (TMB). Absorbance was read at 450 nm.

### Antibody characterization

Kinetic interactions between the antibodies and His-tagged receptor binding domain (RBD, amino acids 319-537) (Acro Biosystems, Newark, NJ) protein was measured at 25°C using BIAcore T200 surface plasmon resonance (SPR) (GE Healthcare). STI-1499 or STI-2020 antibody was covalently immobilized on a CM5 sensor chip to approximately 500 and 100 resonance units (RU), respectively using standard N-hydroxysuccinimide/N-Ethyl-N′-(3-dimethylaminopropyl) carbodiimide hydrochloride (NHS/EDC) coupling methodology. The SARS-CoV-2 RBD protein was coated at five different dilutions (range 0.25?nM–60?nM) in a running buffer of 0.01 M HEPES pH 7.4, 0.15 M NaCl, 3 mM EDTA, 0.05% v/v Surfactant P20 (HBS EP+). The 6.7 nM antigen concentration was measured twice. Blank flow cells were used for correction of the binding response. All measurements were conducted in HBS-EP+ buffer with a flow rate of 30 μl/minute. A 1:1 binding model and was used to fit the data.

### Expression and purification of STI-1499 and STI-2020 monoclonal antibodies

STI-1499 was expressed using a two-vector transient expression protocol. Briefly, 2⨯10^6^/ml CHO-S cells were co-transfected with 1μg of total DNA of the heavy and light chain plasmids, at 1:4 DNA: Polyethyleneimine (PEI) ratio (μg/μg). After 24 hours of incubation with shaking (125 rpm) at 28°C, the culture conditions were adjusted to incubation at 37°C with shaking in a 5% CO_2_ chamber for 10-14 days. STI-1499 was purified from the CHO culture supernatants by affinity chromatography with MabSelectSureLX™ protein A resin using the AktaPure platform.

STI-2020 was produced using a double gene vector stable pool expression protocol. Briefly, the pXC double gene vector (Lonza) encoding both the heavy and light chain genes was used to generate polyclonal pools of stably transfected CHO-S cells, according to manufacturer’s recommended transfection and selection protocols. The stable pool was cultured with shaking (140 rpm) at 37°C in a 5% CO_2_ incubator for 14 days. STI-2020 was purified to homogeneity from the CHO culture supernatants first by affinity chromatography with MabSelectSureLX™ protein A resin and then by ion exchange chromatography using SPImpres resin.

### Cell based Spike binding assay

Mammalian expression vectors were constructed either by cloning of the synthesized gene encoding SARS-CoV-2 G614 Spike protein (UniprotKB, SPIKE-SARS2) or, for SARS-CoV-2 D614 Spike protein, via site-directed mutagenesis of the G614 Spike protein gene. HEK293 cells were transfected using FuGeneHD transfection reagent according to manufacturer’s protocol (Promega, Cat # E2311). 48 hours post-transfection, cells were harvested using enzyme free cell dissociation buffer (ThermoFisher, #13151014.), washed once and resuspended in FACS buffer (DPBS + 2% FBS) at 4⨯10^6^ cells/ml. For antibody binding to the cells expressing the Spike proteins, the cells were dispensed into wells of a 96-well plate (25 μl per well), and an equal volume of 2x final concentration of serially-diluted anti-S1 antibody solution was added. After incubation on ice for 45 minutes, the cells were washed with 150 μl FACS buffer. Detection of bound antibody was carried out by staining the cells with 50 μl of 1:500 diluted APC AffiniPure F(ab’)2 Fragment (Goat Anti-Human IgG (H+L). Jackson ImmunoResearch, Cat# 109-136-4098) for 20 minutes on ice. The cells were washed once with 150 μl FACS buffer and analyzed using flow cytometry. A sigmoidal four-parameter logistic equation was used for fitting the MFI vs. mAb concentration data set to extract EC50 values (GraphPad Prism 8.3.0 software).

### Cells and Viruses

Vero E6 cells were maintained in Dulbecco’s modified Eagle’s medium (DMEM, Corning, NY) supplemented with 10% fetal bovine serum (FBS, Thermo Fisher Scientific, MA), 1% penicillin– streptomycin, and L-glutamine. The P3 stock of the SARS-CoV-2 (USA/WA-1/2020) isolate and the P1 stock of the SARS-CoV-2 (2020001) isolate were obtained from The World Reference Center for Emerging Viruses and Arboviruses (WRCEVA) at the University of Texas Medical Branch. The viruses were propagated in Vero E6 cells and cell culture supernatant of P4 and P2 stocks, respectively, were stored at −80°C under BSL3 conditions.

### SARS-CoV-2 neutralization assay

The day before infection, 2⨯10^4^ Vero E6 cells were plated to 96-well plates and incubated at 37° C, 5% CO_2_. Protein samples (starting from 200 µg/ml) were 2-fold serially diluted in infection media (DMEM+2%FBS) for a total of 8 dilutions. Sixty microliters of diluted samples were incubated with 100 50% tissue culture infective doses (TCID_50_) of SARS-CoV-2 in 60 µl for 1 h at 37°C. One-hundred microliters of the protein/virus mixture were subsequently used to infect monolayers of Vero E6 cells grown on 96-well plates in quadruplicate. Virus supernatant was removed and replaced with fresh medium after 1 h of culture at 37°C. Cytopathic effect (CPE) in each well was observed daily and recorded on day 3 post-infection. At the end of the study, the media was aspirated. Cells were fixed with 10% formalin and stained with 0.25% crystal violet. The neutralizing titers of protein that completely prevented CPE in 50% of the wells (NT_50_) were calculated following the Reed & Muench method^30^.

### Hamster challenge experiments

Male and female Syrian golden hamsters were obtained from Charles River Laboratories at 6 weeks of age. Hamsters were inoculated intranasally (i.n.) with 10^5^ TCID_50_ of SARS-CoV-2 in 100 µl of sterile PBS on day 0. Antibody treatments were administered intravenously (i.v.) with monoclonal antibodies (mAbs) against SARS-CoV-2 Spike, or isotype control mAb in up to 350 µl of sterile PBS at 1 hour-post inoculation. Animals were monitored for illness and mortality for 10 days post-inoculation and clinical observations were recorded daily. Body weights and temperatures were recorded at least once every 48 hours throughout the experiment. On day 5 post-infection, 5 animals from each treatment group were sacrificed and virus titers in the lungs of these animals were measured using a SARS-CoV-2 virus TCID_50_ assay. Average % weight change on each experimental day were compared with the isotype control mAb-treated group using 2-way ANOVA following Dunnett’s multiple comparisons test. All animals were housed in animal biosafety level-2 (ABSL-2) and ABSL-3 facilities in Galveston National Laboratory at the University of Texas Medical Branch. All animal studies were reviewed and approved by the Institutional Animal Care and Use Committee at the University of Texas Medical Branch and were conducted according to the National Institutes of Health guidelines.

### Determination of infectious virus titers in the lung

Prior to initiation of the experiment, five animals from each treatment group (n=10) were designated for virus titration in lung tissue at 5 days post inoculation. On the assigned day, animals were euthanized, lung tissue samples were collected from each animal, and a portion of the tissue (0.08-0.3g) was placed into pre-labeled microcentrifuge tubes containing 5 mm stainless steel beads (Qiagen Inc., CA). Lung samples were homogenized with DMEM + 2% FBS in a TissueLyser (Qiagen Inc., CA) operated at 25-30 Hz for four minutes. Tubes were centrifuged and clarified homogenate was serially diluted 10-fold with DMEM+2% FBS. From material representing each serial dilution step, 100 μl was transferred to each of four wells of a 96-well plated seeded with Vero E6 cells. Plates were then incubated for 72-96 hours at 37° C, 5% CO_2_. Cells were subsequently fixed with 10% formalin and stained with 0.25% crystal violet solution. TCID_50_ values were calculated by the method of Reed and Muench^30^.

## RESULTS

The G-MAB library was panned using a recombinant His-tagged SARS-CoV-2 Spike S1 subunit to allow for selection of candidates with high affinity to the target antigen, as outlined in Figure 1. In summary, following confirmation of S1 binding by ELISA, clonal scFv preparations were tested in a competition ELISA format for disruption of Spike S1:ACE2 binding. Candidate scFvs with high S1 binding affinity and/or the capacity to block Spike S1:ACE2 binding were converted into and expressed as full length human IgG1 antibodies.

**Figure 1.**
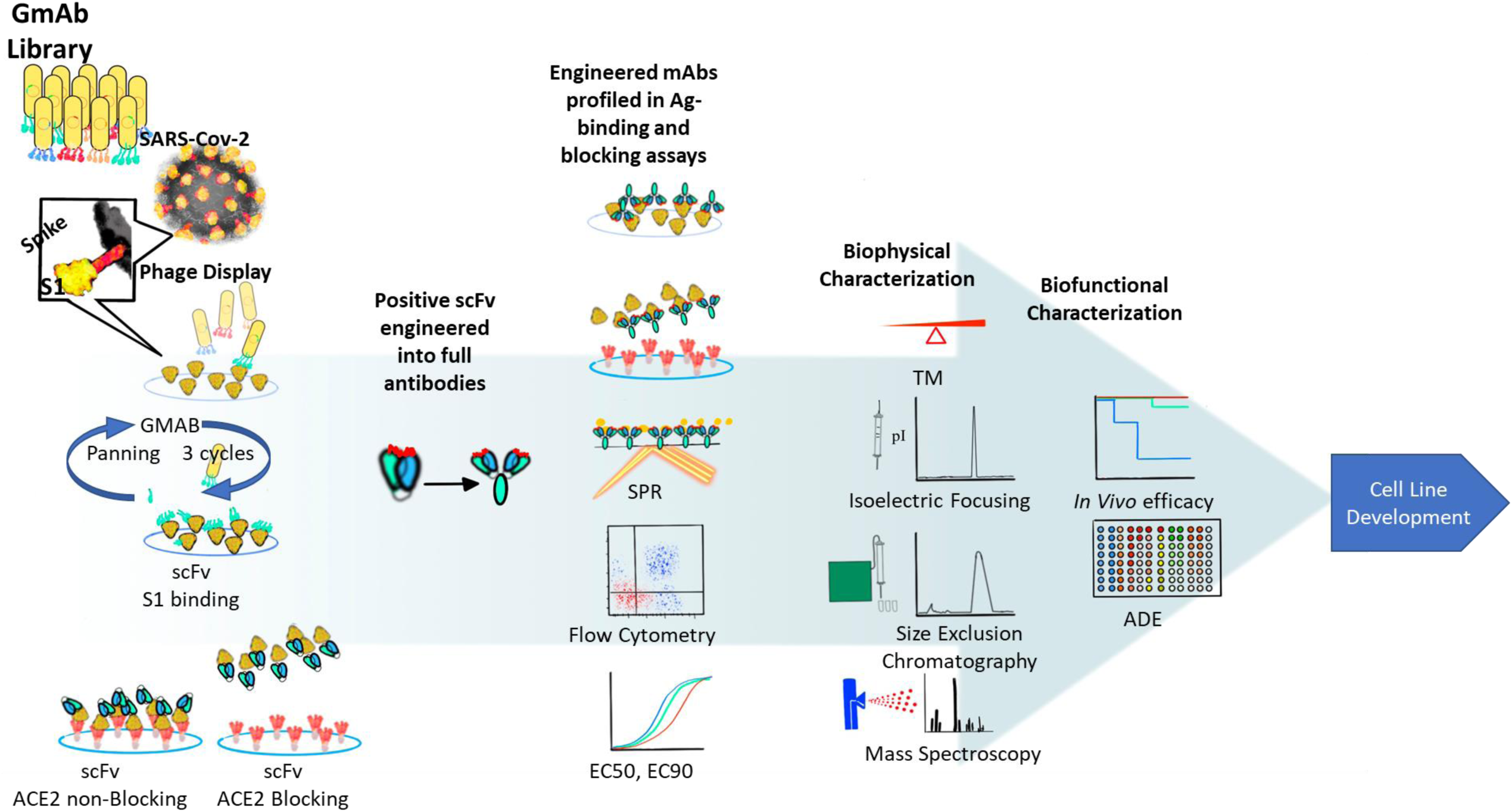
G-MAB SARS-CoV-2 Antibody Discovery Workflow. The G-MAB phage display library was panned for SARS-CoV-2 Spike S1 subunit-binding scFv fragments. Following confirmation of binding activity and blocking of S1:ACE2 interactions by candidate scFvs, the most potent of these candidates were converted to IgG1 antibodies bearing the LALA Fc modification. Candidate nAbs were characterized for binding of Spike S1 subunit and neutralization of related clinical SARS-CoV-2 isolates. Affinity maturation of potent nAbs was carried out in parallel to biophysical profiling, cell line development, and evaluation of protective efficacy for the parental nAb, STI-1499.

In an effort to mitigate the risk of ADE resulting from administration of G-MAB-derived nAbs, the IgG1 Fc region were modified by introducing specific amino acid substitutions (L234A, L235A [LALA]) ^31^. The LALA Fc modification reduces binding affinity to the Fcγ receptors, thus addressing the potential ADE risks associated with delivery of unmodified recombinant IgG1 neutralizing antibody or convalescent sera to COVID-19 patients, while providing a similar blockade to interactions between SARS-COV-2 and the angiotensin-converting enzyme 2 (ACE2) receptor expressed on susceptible cells in the lung and gut epithelia ^32, 33, 34^. Partitioning of virus away from susceptible cells in this manner may allow for clearance of nAb-bound virus and a corresponding decrease in the pace and degree of COVID-19 pathogenesis in treated individuals.

### Functional profiling and antibody affinity maturation

The nAb candidate termed STI-1499 displayed potent SARS-CoV-2 neutralizing activity superior to that of other G-MAB nAb hits, with an IC_50_ in the virus neutralization assay of 3.2 μg/ml (Figure 2A). In parallel to STI-1499 preclinical profiling and cell line development efforts, affinity maturation of STI-1499 was undertaken, with systematic amino acid changes engineered within complementarity determination regions (CDRs) of the STI-1499 heavy and light chains. For individual CDR regions from STI-1499, a library of variants was screened for binding to the Spike S1 subunit by ELISA (Figure 2B). Variants bearing combinations of affinity-enhancing CDR amino acid changes were then systematically engineered and expressed as IgG1 LALA antibodies. Based on Spike S1 ELISA binding results, the most potent clone of the single CDR and combination CDR variants, STI-2020, was further profiled for binding affinity using surface plasmon resonance (SPR). STI-2020 bound to the RBD region of the Spike protein with an affinity of 1.3 nM, a 35-fold affinity improvement over that of the parental STI-1499 (Figure 2C).

**Figure 2.**
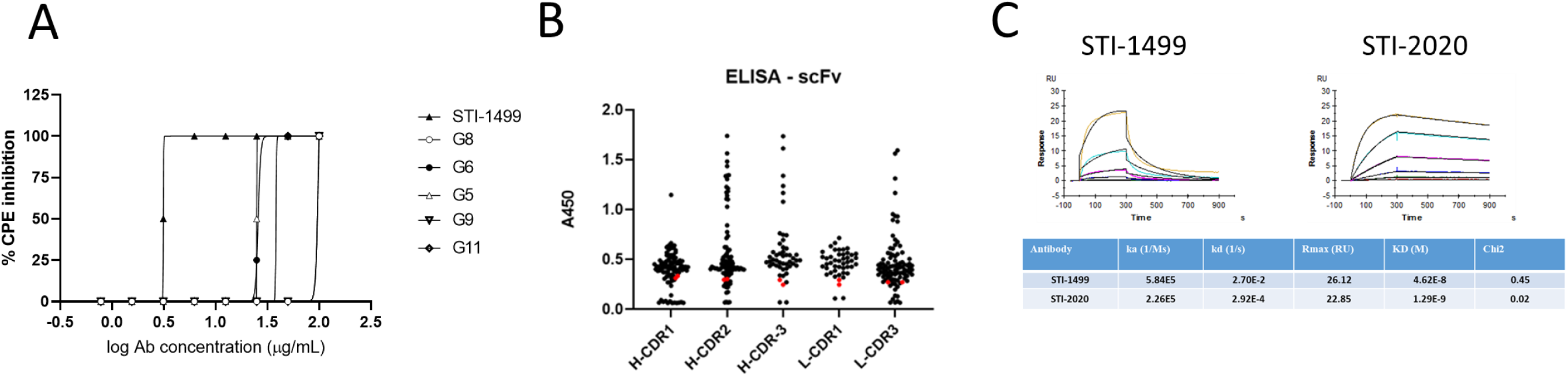
Identification of STI-1499 nAb and Affinity Maturation to Identify STI-2020. **(A)** A panel of candidate nAbs was evaluated in a SARS-CoV-2 neutralization assay (USA/WA-1/2020 isolate). The STI-1499 (closed triangle) demonstrated potent virus neutralization with a calculated IC_50_ of 3.13 μg/ml. **(B)** Five independent scFv phage libraries covering HCDR1-3, LCDR1, and LCDR3 were generated and screened. 94 single clones (black dots) were randomly selected and together with 2 clones representing wild type STI-1499 as reference (red dots) were analyzed by ELISA. **(C)**. Affinity measurement of STI-1499 and STI-2020. The antibody affinities were measured using SPR on a BIAcore T200 instrument.

### Spectrum of activity of SARS-CoV-2 neutralizing antibodies

STI-1499 and STI-2020 were tested for binding to full length Spike proteins derived from naturally emerging viruses including the USA/WA-1/2020 (WA-1) isolate as well as an emerging SARS-CoV-2 isolate that bears a D →G mutation at amino acid residue 614 in the Spike protein (D614G). Binding studies demonstrated equivalent binding affinity for both nAbs against cell surface-expressed Spike protein with the D614G mutation as well as that of the WA-1 isolate (Figure 3A).

**Figure 3.**
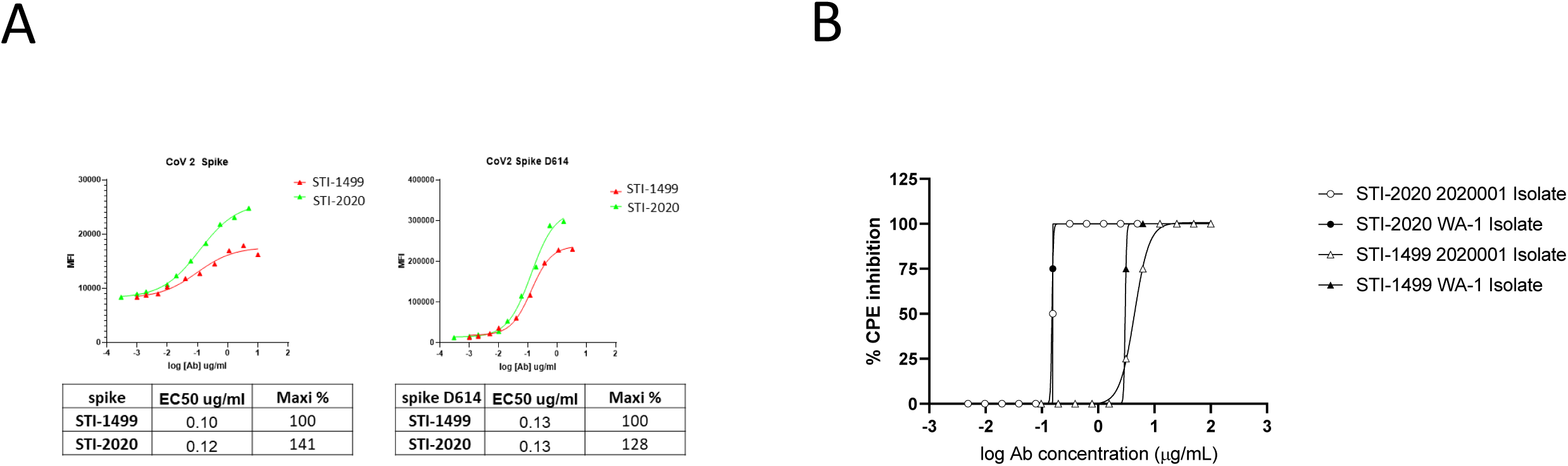
Binding of Spike Protein and Neutralization of Emerging Pandemic Isolate by STI-1499 and STI-2020. **(A)** Spike protein derived from the WA-1 and 2020001 (D614G) SARS-CoV-2 isolates were independently expressed on the surface of HEK 293 cells. Serially-diluted STI-1499 or STI-2020 were assayed for Spike protein binding by flow cytometry. To quantify antibody binding, mean fluorescence intensity was measured for each dilution tested and the EC_50_ value was calculated for each nAb. **(B)** STI-1499 and STI-2020 were evaluated in a SARS-CoV-2 neutralization assay for potency against the USA/WA-1/2020 isolate and the 2020001 (D614G) isolate.

To determine the neutralizing activity of STI-1499 and STI-2020 against the aforementioned SARS-CoV-2 clinical isolates (*vide supra*), nAbs were evaluated in the virus neutralization assay. Both nAbs exhibited equipotent neutralization of the virus isolates. In keeping with observed improved binding properties, STI-2020 neutralized both isolates with an IC_50_ of 0.055 μg/ml and an IC_99_ of 0.078 μg/ml, a greater than 50-fold enhancement of neutralizing potency over the parental STI-1499 nAb (Figure 3B).

### Treatment with STI-1499 or STI-2020 in a hamster model of COVID-19

SARS-CoV-2 pathogenesis in the Syrian golden hamster model of infection provides a means of assessing nAb activity in a preclinical model of respiratory disease. Animals inoculated with 1×10^5^ TCID_50_ of SARS-CoV-2 intranasally were treated with either STI-1499 (Figure 4) or STI-2020 (Figure 5) administered intravenously (IV) at 1 hour post-infection. Weight change as a percentage of starting weight was recorded and graphed for each animal (Figures 4 Panel A, Figure 5 Panel A). Animals in the STI-1499 treatment groups were administered a single dose of either 50, 500, or 2000 μg IV. To control for effects of the virus infection on animal growth, uninfected animals (UI) were administered 2000 μg of the control IgG1 (IsoCtl) IV. Animals in this group displayed no clinical signs and gained weight throughout the course of the experiment. Hamsters infected with SARS-CoV-2 and administered 2000 μg of IsoCtl experienced steady weight loss over the first five days of infection, with maximal average weight loss % in this group reaching −5.7% ± 3.3 at 5 days-post-infection (d.p.i.). The duration and degree of weight loss in the 50 μg and 500 μg STI-1499 treatment groups were similar to that of animals administered IsoCtl mAb. Following administration of 2000 μg STI-1499, animals experienced transient minor weight loss over the first two days post-infection followed by steady weight gain on subsequent days. By day 4 d.p.i., and continuing on until the end of the experiment on 10 d.p.i., the effects on SARS-CoV-2 infection on the average % weight change of animals treated with 2000 μg STI-1499 was significantly different than that in animals treated with control IgG (Figure 4, Panel C). Animals treated with 2000 μg STI-1499 began to gain weight by day 3 d.p.i. and had reached an average weight gain of 5.8% ± 4.1 by 10 d.p.i. while animals treated with IsoCtl mAb continued to lose weight until 5 d.p.i. and, on average, only returned to day 0 weight levels by 10 d.p.i.. At 5 d.p.i., lungs from 5 out of 10 animals (designated prior to initiation of the experiment) from each experimental group were harvested and virus titers were quantified as TCID_50_/g. Treatment with 2000 μg STI-1499 resulted in an average lung titer of 1.38 x 10^4^ TCID_50_/g of tissue, an approximately 27-fold reduction compared to average lung titers in IsoCtl-treated animals. Notably, lung titers for three of the five animals from the 2000 μg STI-1499 group were below the limit of detection (<3.6×10^2^ TCID_50_/g). The remaining two animals displayed lung titers approximately 10-fold below that of the IsoCtl-treated group. Consistent with a partial protective effect following a dose of 500 μg of STI-1499, the average TCID_50_/g of lung tissue among animals in this treatment group were reduced 2.5-fold as compared to the IsoCtl treatment group.

**Figure 4.**
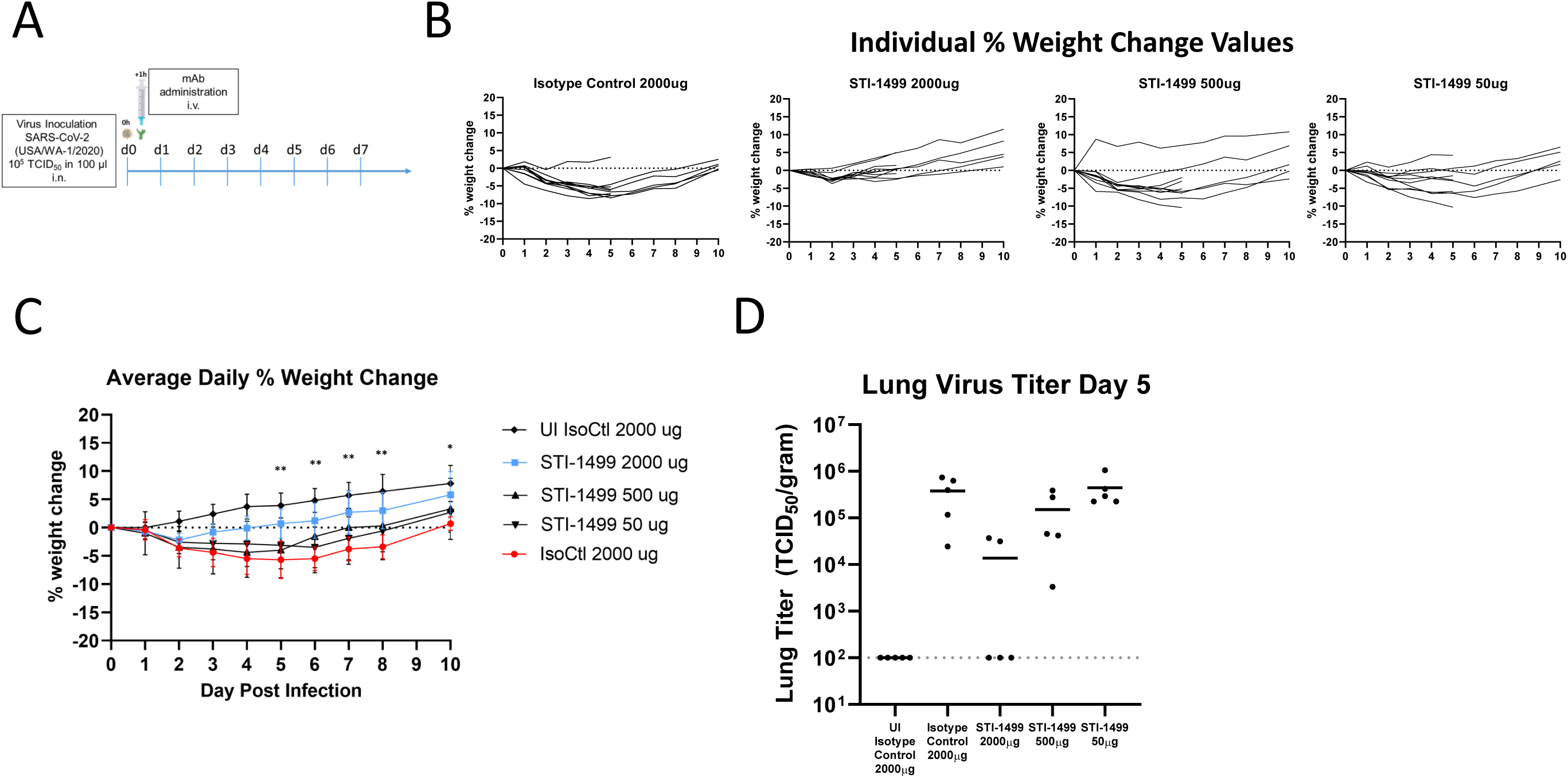
Protective Efficacy of STI-1499 in the Syrian Golden Hamster Model of COVID-19. **(A)** Male hamsters were inoculated with SARS-CoV-2 WA-1 isolate on day 0. One-hour post-infection, animals (n=10 per group) were administered a single intravenous dose of Control IgG (2000 μg) or STI-1499 (50 μg, 500 μg, or 2000 μg). Daily weight changes from day 0 to day 10 were recorded and **(B)** plotted for each individual animal. **(C)** Average % daily weight change ± standard deviation was plotted for each group. Days on which there was a significant difference in average % weight change between STI-1499 2000 μg-treated animals and Control IgG 2000 μg-treated animals are denoted by *(p-values < 0.05) or **(p-values <0.01). **(D)** Lung tissues collected from five animals per group and virus titers were determined on day 5.

**Figure 5.**
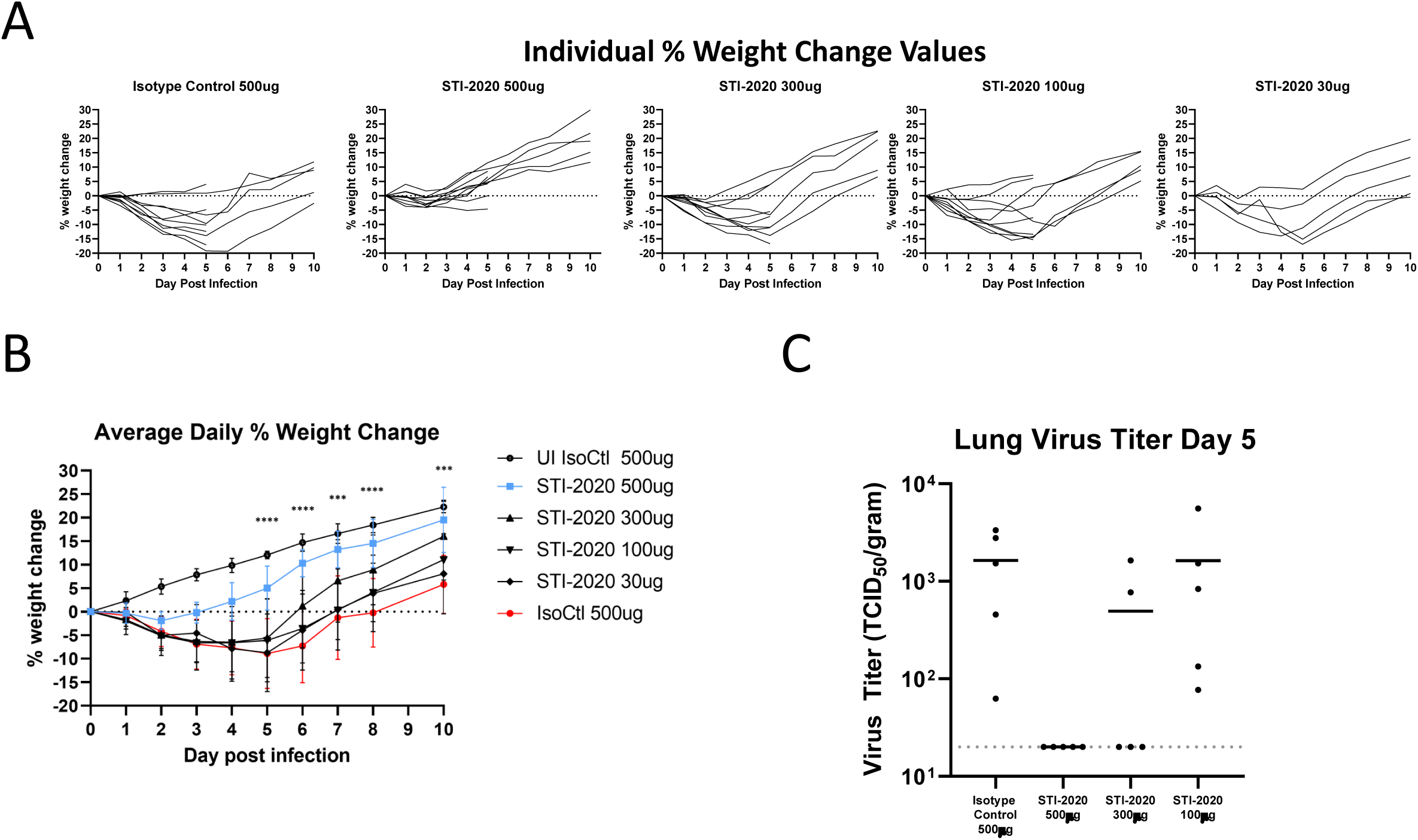
Protective Efficacy of STI-2020 in the Syrian Golden Hamster Model of COVID-19. Female hamsters were inoculated with SARS-CoV-2 WA-1 isolate on day 0. One-hour post-infection, animals (n=10 per group) were administered a single intravenous dose of Control IgG (500 μg) or STI-2020 (30 μg, 100 μg, 300 μg, or 500 μg). Daily weight changes from day 0 to day 10 were recorded and **(A)** plotted for each individual animal. **(B)** Average % daily weight change ± standard deviation was plotted for each group. Days on which there was a significant difference in average % weight change between STI-2020 500 μg-treated animals and Control IgG 500 μg-treated animals are denoted by ***(p-values ≤ 0.0003) or ****(p-values ≤ 0.0001). (D) Lung tissues collected from five animals per group and virus titers were determined on day 5.

Animals in the STI-2020 treatment groups were administered a single dose of either 30, 100, 300, or 500 μg IV. Administration of a 500 μg dose of STI-2020 resulted in maximum average percentage body weight loss of −1.9%, which occurred on 2 d.p.i.. After that, animals treated with 500 ug STI-2020 maintained an average body weight that as a percentage of day 0 weight was significantly different on days 4, 5, 6, 7, 8, and 10 than the average weight measured among IsoCtl-treated animals. As with STI-1499-treated hamsters, virus titers in the lung were measured in 5 of 10 animals from each treatment group. In IsoCtl-treated animals, an average of 1.6×10^3^ TCID50/g of lung tissue was detected. Treatment with 500 μg STI-2020 resulted in reduction of lung titers below the level of detection in all animals tested, a STI-2020 - treatment-related lung titer reduction of 80-fold at minimum. In the 300 μg STI-2020 treatment group, the average lung titer was reduced below the level of detection in 3 of 5 animals tested, while 2 of 5 animals had lung titers of similar magnitude to those measured in IsoCtl-treated animals. Animals with undetectable lung virus titers in the 300 μg treatment group also experienced only moderate weight loss compared to animals with detectable lung virus titers in this group (d5 values of −11.1% and −16.7%) No changes in average lung titer compared to IsoCtl-treatment were detected in animals from the 100 μg STI-2020 treatment group.

In summary, STI-1499 demonstrated promising protective efficacy in the hamster model at a dose of 2000 μg. The observed overall enhancement of *in vitro* potency for the affinity matured nAb, STI-2020, translated into increased protection against disease in the hamster challenge model, with a greater than 4-fold increase in protective efficacy over that of STI-1499.

## DISCUSSION

In this study, we detail the initial discovery and profiling of a SARS-CoV-2 nAb isolated from a phage display antibody library derived from the B-cell repertoire of over 600 healthy normal individuals. This approach complements the isolation of nAbs collected from natural sources, such as convalescent patients or vaccinated healthy individuals, and allows for isolation of nAbs recognizing neutralizing epitopes developed outside the temporal and biological context of the pathogen/antigen-specific immune response. The parental nAb STI-1499 and the affinity-matured derivative STI-2020 were characterized for their biochemical and function properties *in vitro* and *in vivo* against prevalent clinical isolates of SARS CoV-2. Both nAbs demonstrated potent functionality in these assays, which is strongly supportive of continued preclinical and clinical development. Notably, demonstration of improvement in neutralizing potency *in vitro* for STI-2020 over that of STI-1499 was predictive of increased protective efficacy in the hamster challenge model following post-exposure therapeutic treatment. In addition, *in vivo* efficacy was achieved in the presence of the ADE-mitigating LALA Fc modification, providing further confidence that this modification might add to the utility of nAbs under clinical evaluation for use in the treatment of COVID-19. Studies aimed at establishing the therapeutic and prophylactic treatment window in the hamster model as well as demonstrating efficacy following administration of nAb proteins by non-parenteral routes are currently underway.

In addition, introduction of nAb-encoding DNA plasmids provides a low cost, long-acting means of establishing antibody-mediated immunity, and experiments in rodents are ongoing and will determine the suitability of STI-2020 for use in this context. Treatments using STI-2020 nAb protein or DNA-encoding of nAbs in combination with other identified nAbs targeting non-overlapping epitopes on the Spike S1 subunit to constitute a nAb cocktail approach will provide a means of increasing the barrier to treatment resistance as the prevalence of naturally emerging and treatment-associated virus variants increases. This approach is under investigation and in preclinical development.

As emergence of novel pathogens will continue to pose a significant threat to humanity, the rapid discovery of potent nAbs against these yet-unknown microbes will remain paramount to global health efforts. The discovery and development of anti-SARS-CoV-2 nAbs STI-1499 and STI-2020 in less than 6 months without the need of samples from infected individuals could serve as blueprint for future rapid response efforts.

Panning of the G-MAB library consisting of fully human antibody sequences allowed for the expeditious progression of lead nAb candidates from initial screening to the production cell line generation stage, avoiding additional development steps such as subsequent humanization of the identified antibodies and associated immunogenicity concerns of non-human antibody-based therapeutics. This has enabled an efficient and expedited discovery and development of SARS-CoV-2 nAbs starting in March 2020, thus taking less 6 months from initiation of the project to completed IND filing. Notably, STI-1499 has recently received FDA clearance to commence first-in-human studies making it the first clinical-stage anti-SARS-CoV-2 nAb derived not from convalescent patients but from a fully human phage display antibody library, and it is anticipated that dosing of the first patients will begin in October 2020.

## Author Contributions

R.A., H.J., S.P., M.B., H.Z., Y.Z., J.M. conceptualized and designed experiments. J.M., B.B., X.C., N.D., S.J., L.K., K.L., D.L., L.L., J.T.M., K.M., C.P., S.R., R.S., Y.Z., H.Z performed experiments and J.M., X.C., L.K., S.P., Y.Z., H.Z., R.A. analyzed data. R.A., H.J., J.M., S.P. wrote the paper.

## Competing interests

Sorrento authors own options and/or stock of the company. This work has been described in one or more provisional patent applications. HJ is an officer at Sorrento Therapeutics, Inc..

